# Development of a preclinical screening platform for clinically relevant therapy of Dravet syndrome

**DOI:** 10.1101/2024.08.13.607806

**Authors:** Jeffrey A. Mensah, Kyle E. Thomson, Jennifer L. Huff, Tia Freeman, Christopher A. Reilly, Joseph E. Rower, Cameron S. Metcalf, Karen S. Wilcox

## Abstract

**Background:** Patients with drug-resistant epilepsy, including Dravet syndrome (DS), are frequently prescribed multiple antiseizure medications (ASMs). Nevertheless, people with DS often have inadequate seizure control, and there is an ongoing unmet clinical need to identify novel therapeutics. As a proof-of-principle study to further validate and characterize the *Scn1a*^*A1783V/WT*^ mouse model and identify a drug screening paradigm with face, construct, and predictive value, we assessed the efficacy of subchronic administration of stiripentol add-on to clobazam and valproic acid at clinically relevant doses using the *Scn1a*^*A1783V/WT*^ mouse model.

**Methods:** We evaluated the efficacy of STP add-on to CLB and VPA using hyperthermia-induced and video-EEG monitoring of spontaneous seizure tests following a 14-day treatment. VPA was delivered via osmotic minipump, while STP and CLB were administered via food pellets delivered through automatic feeders. Bioanalytical assays were performed to evaluate drug concentrations in plasma and brain using liquid chromatography-tandem mass spectrometry.

**Results:** STP, CLB, N-desmethylclobazam, and VPA all yielded plasma concentrations within the human therapeutic plasma concentrations range. STP added to CLB and VPA significantly elevated the seizing temperatures in the hyperthermia-induced seizure assay. CLB, VPA, and STP coadministration significantly reduced spontaneous seizure frequency compared to CLB and VPA combined.

**Significance:** This research lays the groundwork for exploring effective add-on compounds to CLB and VPA in treating DS. The study further highlights the utility of the *Scn1a*^*A1783V/WT*^ mice in discovering therapies for DS-associated pharmacoresistant seizures.

**Key points:**

- Integrating pharmacokinetic studies to guide the selection of doses in preclinical studies to achieve target concentrations comparable to the human therapeutic range is crucial in successfully translating animal drug development studies to clinical use.
- STP add-on to CLB and VPA significantly reduced spontaneous seizure frequency in *Scn1a*^*A1783V/WT*^ mice.
- A triple-drug polytherapy approach that mimics the clinical treatment paradigm will be an essential preclinical drug screening strategy for identifying novel investigational compounds for Dravet syndrome.

## Introduction

Dravet syndrome (DS) is a severe, early-onset genetic epileptic disorder [1]. Prolonged febrile, recurrent spontaneous seizures, and severe intellectual decline mark most cases [2, 3]. Additionally, children with DS are at an estimated 15-fold greater risk of SUDEP than other childhood epilepsies [4]. Despite the availability of several antiseizure medications (ASMs), approximately 30% of patients with DS remain pharmacoresistant [5]. Identifying effective drugs and treatment paradigms is crucial for achieving adequate seizure control for DS; thus, drug screening models with good face, construct, and predictive validity are essential in our search for efficacious ASMs.

Preclinical and clinical drug development of newer effective ASMs for DS has received significant attention. Considering that several ASMs often do not progress from the preclinical development phase to clinical trials, the use of appropriate genetic models of DS that better recapitulate the crucial aspects of the disorder, such as recurrent spontaneous and febrile seizures like in humans, is imperative for successful therapy discovery [6]. Additionally, considering that DS is a chronic brain disorder and those with DS are at high risk of SUDEP [7], a relevant drug screening strategy, such as subchronic treatment paradigms, that mirror clinical response more closely is critical. Thus, the combined use of an effective drug screening paradigm and an etiologically relevant animal model of DS provides an essential platform for identifying efficacious DS therapeutics.

We previously demonstrated [8] that the acute hyperthermia-induced seizure test in *Scn1a*^*A1783V/WT*^ mice was a pharmacoresistant assay and could identify novel efficacious therapies for DS. We further showed that acute intraperitoneal injections of stiripentol (STP) in combination with clobazam (CLB) and sodium valproate (VPA) were effective in increasing the temperature at which hyperthermia-induced seizures occurred in the *Scn1a*^*A1783V/WT*^ mouse model of DS [9]. This study showed that a triple-drug therapy paradigm that mimics clinical standards for add-on therapies in DS is a useful preclinical strategy for discovering novel investigational compounds for DS. Using pharmacokinetic (PK) analysis of samples collected following single injections of each drug, we found that brain levels of CLB and VPA metabolites were elevated following STP administration [9]. This observation may explain the potentiated efficacy of triple-drug therapy. However, whether this triple-drug therapy is effective against spontaneous recurrent seizures (SRSs) at doses sufficient to provide steady-state therapeutic plasma concentrations has yet to be tested.

The NINDS Epilepsy Therapy Screening Program (ETSP) aims to facilitate the discovery of new efficacious therapeutics and enhance translation using relevant models and screening approaches that best mimic clinical data [10]. In the present study, we designed an effective subchronic treatment paradigm of a clinically relevant second-line, multi-drug combination therapy for DS. PK studies were utilized to ascertain the efficacy of drug doses that achieved plasma concentrations within the human plasma therapeutic range. Subchronic (14-day) treatment of STP, when given in combination with CLB and VPA, significantly increased the temperature at which *Scn1a*^*A1783V/WT*^ mice seized. Additionally, CLB+VPA+STP treatment significantly reduced spontaneous seizures in DS mice and conferred seizure freedom in a subset of mice. Our findings align with what is typically observed in clinical practice and provide important validation of such preclinical polypharmacy screening approaches. The current study demonstrates the practicality of using both spontaneous recurrent seizures and hyperthermia-induced acute seizure phenotype in the *Scn1a*^*A1783V/WT*^ mouse model of DS as a useful preclinical platform for designing and evaluating subchronic treatment strategy for clinically relevant combination therapies in DS. Our future studies will evaluate investigational compounds in combination with VPA and CLB to identify new therapies for people with drug-refractory DS.

## Method

### Animal

The University of Utah Institutional Animal Care and Use Committee approved all animal care and experimental procedures. Animal experiments were conducted per Animal Research: Reporting of In Vivo Experiments Guidelines (https://www.nc3rs.org.uk/arrive-guidelines). Experimental mice were generated, as reported by Pernici et al. [8]. Both female and male heterozygous and age-matched wild-type littermates were used for experiments. Mice were group-housed in a pathogen-free mouse facility under standard laboratory conditions (14-h light/10-h dark cycle) and had access to food and water ad libitum, except during hyperthermia-induced seizure experiments when they were transferred to the experimental room approximately 60 mins before testing. A case report form was used to record a detailed list of common data elements and confirm all study aspects.

### Materials

CLB and VPA were purchased as powders from Sigma-Aldrich. N desmethylclobazam (N-CLB) and VPA-^13^C_6_were purchased as 1.0 mg/mL stock solutions in MeOH, while CLB-^13^C_6_ and NCLB-^13^C_6_ were purchased as 0.1 mg/mL stock solutions in acetonitrile from Sigma Aldrich (city, state). STP and STP-d_9_ were purchased as powders from Toronto Research Chemicals (Toronto, Canada)

### Drug-in-food pellet formulation

The appropriate amount of experimental drug(s) was weighed and placed in a large glass mortar container. The required powder (Bioserv rodent MD’s pre-mix, chocolate flavor, placebo powder, product # F7762) (Flemington, NJ) was also weighed into a Tupperware/container. We then performed geometric addition of drug(s) and chow powder using a large glass mortar and pestle until all materials were mixed. The resulting mixture was transferred into a KitchenAid mixer for approximately 10 minutes at low speed. With the mixer on, water is gradually sprayed into the mixer, followed by a few rotations until the required volume of water is added. The mixture is then granulated 1x using a 1.6 mm Erweka Machine sieve. The granules were then evenly spread out and dried for approximately 1 hour. Materials were then transferred to Tupperware and pressed into pellets. Food pellets were left in a hood to dry overnight.

### Drug preparation and administration

The current study’s CLB, VPA, and STP dosage regimen was selected based on PK data reported by our group [9]. CLB and STP were formulated into chow using the Bioserv pre-mix powder as described. CLB (250 mg/kg/day food) and STP (4000 mg/kg/day food) were given to mice via pellets at 4-hour intervals using an automated drug-in-pellet delivery system [11]. The estimated doses of CLB and STP were based on a 20 g mouse and an average food consumption of 3 g/day. VPA was administered by subcutaneous sub-chronic (14-day) administration using the Alzet osmotic minipump. VPA (1.5 g/mL at 0.5 μL/hr flow rate) was prepared in 0.9% NaCl (Table 1).

**Table 1:** Plasma and brain concentrations of CLB, N-CLB, VPA, and STP following a 14-day treatment in adult *Scn1a*^*A1783V/WT*^ mice.

| Compound Measured<br>(treatment paradigm) | Drug Dose | Mean concentration following<br>a 14-day treatment |  | Human therapeutic<br>plasma<br>concentration<br>range |
| --- | --- | --- | --- | --- |
| | | Plasma<br>concentration<br>($\mu\text{g/mL}$) | Brain<br>concentration<br>($\mu\text{g/g}$) | |
| **CLB** (CLB + VPA) | 250 mg/kg/day food<br>pellet | 0.03 | 0.025 | 0.03 - 0.3 $\mu\text{g/mL}$<sup>[29]</sup> |
| **CLB** (CLB + VPA + STP) | 76 mg/kg/day food<br>pellet | 0.07 | 0.011 |  |
| **N-CLB** (CLB + VPA) | | 0.8 | 1.75 | 0.3 - 3 $\mu\text{g/mL}$<sup>[29]</sup> |
| **N-CLB** (CLB + VPA + STP) |  | 3.2 | 7.15 |  |
| **VPA** (CLB + VPA) | 1.5 g/mL at 0.5 $\mu\text{L/hr}$<br>flow rate | 84.3 | 14.9 | 50 - 100 $\mu\text{g/mL}$<sup>[30]</sup> |
| **VPA** (CLB + VPA + STP) |  | 96.9 | 21.25 |  |
| **STP** | 4000 mg/kg/day food<br>pellet | 17.33 | 1.05 | 4 -22 $\mu\text{g/mL}$<sup>[31]</sup> |

Abbreviations: Conc, concentration; CLB, clobazam; N-CLB, N-desmethyl clobazam; STP, stiripentol, VPA, sodium valproate
Drug doses were calculated using PK data from Mensah et al. [9].
n= 5-7 mice per group

### Osmotic mini-pump implantation surgery

Minipumps (Alzet model 2002) were filled with VPA and submerged in saline overnight. Briefly, mice were anesthetized with isoflurane (2-4%) and then transferred to a stereotaxic frame equipped with a nose cone for isoflurane delivery. A small incision was made between the scapulae, followed by clearance of the subcutaneous fascia with forceps. Minipumps were placed in the subcutaneous space, and the incision was sutured with surgical silk. Mice were postoperatively administered buprenorphine (0.015 mg/kg, s.c., MWI Animal Health, Boise, ID) for analgesia and enrofloxacin antibiotic (5 mg/kg BW, s.c.) to prevent infection and monitored daily for the entire 14-day treatment period.

### Hyperthermia-induced seizure test

We evaluated the temperature at which *Scn1a*^*A1783V/WT*^ mice seized on Day 0 (pre-treatment), Day 7, and Day 14 of drug treatment. Five to seven-week-old *Scn1a*^*A1783V/WT*^ mice were put in a glass chamber and placed under a heat lamp, and the core temperature was gradually elevated by 1°C every 2 min until a seizure was observed or the temperature reached 42.5°C [12, 13]. Before a test, mice were acclimated to the temperature probe for 5 minutes in a glass chamber. Hyperthermia test was conducted approximately 2 hours following delivery of food pellets with drug via auto feeders to allow enough time to consume all pellets. Body temperature was monitored using a neonate rectal probe (Braintree Scientific, Inc, Braintree, MA) coupled to a TCAT-2LV controller (Physitemp Instruments, Inc, Clifton, NJ). The temperature at which mice seized was recorded. Male and female mice were randomly assigned to treatment groups on Day 0, and the same groups were maintained throughout the study. Group One (n=7) received STP (in food pellets)+CLB (in food pellets)+VPA (s.c., minipump), group 2 (n=7) received CLB (in food pellets)+VPA (s.c., minipump), and group 3 (n=7) received Bioserv Custom food pellets without any test drug + saline (s.c., minipump) (vehicle). Experimenters were blinded to the treatment.

### Bioanalytical study

PK studies of CLB, VPA, and STP to establish doses in *Scn1a*^*A1783V/WT*^ mice that resulted in therapeutic plasma drug concentrations were performed as described by Mensah et al. [9] and is described briefly below.

#### CLB, N-CLB, and VPA

CLB and VPA were diluted in 50:50 methanol:water (MeOH:H_2_O) to 1.0 mg/mL and 10 mg/mL concentrations, respectively. NCLB and VPA-^13^C_6_ were purchased as 1.0 mg/mL stock solutions in MeOH, while CLB-^13^C_6_ and NCLB-^13^C_6_ were purchased as 0.1 mg/mL stock solutions in acetonitrile, all from Sigma Aldrich. Mouse plasma samples were diluted 20-fold before extraction, while mouse brain homogenate was analyzed without dilution. Human plasma and mouse brain homogenate were used as the control matrix for calibrators and quality control (QC) samples for mouse plasma and brain samples, respectively.

A 100 μL aliquot of the 20-fold diluted plasma or undiluted brain homogenate was assayed using a protein precipitation extraction approach. A 50 μL volume of internal standard (IS) (400 pg/μL VPA-^13^C_6_, 40 pg/μL CLB-^13^C_6,_ and NCLB-^13^C_6_ in 10:90 MeOH:H_2_O) was added to each sample aliquot, followed by 1 mL of cold acetonitrile. The samples were vortex-mixed and centrifuged to separate the supernatant from the protein pellet. The supernatant was then transferred to a fresh microcentrifuge tube. The supernatant was centrifuged again to ensure the removal of all solid particulates. The supernatant was transferred into a polypropylene tube and dried under house air (15 psi) in a TurboVap set (40°C). Samples were then reconstituted with 200 μL of 10:90 MeOH:H_2_O, centrifuged, vortex mixed, and transferred to autosampler vials for analysis.

A volume of 10 μL of sample was injected on a Waters Acquity UPLC with a Quattro Premier XE UHPLC-MS/MS system for analysis. Chromatography was performed using a Phenomenex (Torrance, CA) Luna Omega Polar C18, 1.6 μm (2.1 × 50 mm), and gradient elution was maintained at a 300 μL/min flow rate. Mobile phases consisted of (A) 10mM ammonium acetate (pH 5.5) and (B) methanol. Analytes were monitored using the following mass transitions (collision energy, CE): 287.1→245.1 (NCLB, CE=20 V); 293.1→251.1 (NCLB-^13^C_6_, CE=20 V); 301.1→259.1 (CLB, CE=20 V); 307.1→265.1 (CLB-^13^C_6_, CE=20 V); 143.1→143.1 (VPA, CE=5 V); and 149.1→149.1 (VPA-^13^C_6_, CE=5 V). CLB, NCLB, and the IS utilized positive electrospray ionization (ESI), whereas VPA and its internal standard (IS) used negative ESI. The calibration curves utilized 1/x^2^ weighted linear regression between 4-200 ng/mL for CLB and NCLB and 0.2-10 μg/mL for VPA.

#### STP

STP was diluted in 90:10 MeOH:H_2_O to a 1.0 mg/mL concentration. Mouse plasma samples and brain homogenates were diluted 100-fold and 10-fold in 50:50 MeOH:H_2_O. A 200 μL aliquot of the 100-fold diluted plasma sample was analyzed with calibrators, and QC was prepared in 1% mouse plasma in 50:50 MeOH:H_2_O. A 200 μL aliquot of the 10-fold brain homogenate was assayed against calibrators and QCs prepared in 10% mouse brain homogenate in 50:50 MeOH:H_2_O.

A basified liquid-liquid extraction procedure was used to extract STP. A 50 μL volume of 40 pg/μL STP-d_9_ in 50:50 MeOH:H_2_O was added to each sample in polypropylene tubes. 0.5 mL of 5% (v:v) ammonium hydroxide in type-1 water was used to adjust the sample pH and extracted with 2.0 mL of methyl tert-butyl ether (MTBE). The sample was mixed well and centrifuged to separate the organic and aqueous layers. The sample was frozen at −80°C for 30 min, and the organic layer was decanted into a fresh tube. The organic layer was dried under house air (15 psi) in a TurboVap set (40°C). The sample was then reconstituted with 200 μL of 50:50 MeOH:H_2_O and transferred to an autosampler vial for analysis.

A 20 μL sample volume was injected onto a Waters Acquity UPLC and autosampler coupled with a ThermoScientific TSQ Quantum Access MS/MS. Chromatography utilized a Phenomenex Luna Omega PS C18 3μm (2.1 × 100 mm) column, and gradient elution was maintained at a 300 μL/min flow rate. Mobile phases comprised (A) 0.1% formic acid in type-1 water and (B) methanol. STP and the IS were ionized using positive ESI, and the mass transitions (CE) 217.2→187.2 (11 V) and 226.2→196.2 (11 V) were monitored for STP and STP-d_9_, respectively. The assay’s dynamic range was 1-1000 ng/mL, fitting with a 1/x weighted linear regression.

### Electrode implantation surgery

Male and female *Scn1a*^*A1783V/WT*^ mice (8 weeks old) were surgically implanted with a cortical, bipolar electrode (MS333/6-B/SPC, PlasticsOne, Roanoke, VA). Mice were anesthetized with isoflurane (4%) and placed on a stereotaxic frame. Isoflurane (2 %) delivery was maintained for the duration of the surgery. A hole into the skull at AP = −2.3 mm; ML = 2.55 (relative to bregma) was drilled, and a bipolar electrode was placed on the surface of the dura mater, with a ground electrode placed over the cerebellum. Three anchoring screws were placed over the cortex, one contralateral and two ipsilateral of the electrode implantations sites. A dental acrylic (Bosworth Company, Midland, TX) secured the electrode and anchor screws to the skull. Mice were postoperatively administered buprenorphine (0.015 mg/kg, s.c., single administration post-surgery, MWI Animal Health, Boise, ID) for analgesia and enrofloxacin (5 mg/kg BW, s.c.) to prevent infection.

### Video-EEG monitoring and seizure analysis

Mice were tethered for 24hr/day, 7 day/week video-EEG recording after a 14-day recovery from the electrode implantation surgery. We tethered mice to a rotating commutator (Plastics One, Roanoke, VA) connected to an EEG100c amplifier (BIOPAC Systems, Goleta, GA). A BIOPAC MP150 Recording System (BIOPAC Systems, Goleta, GA) was used to digitize EEG signals, and DVP 7020BE Capture cards captured video. Baseline video-EEG recordings were taken for 14 days. During baseline recordings, mice were fed with regular Bioserv pellets (without experimental drugs) via auto feeders. Male and female mice were randomly and proportionally put into treatment groups. One group (n=13) received STP (in food pellets)+CLB (in food pellets)+VPA (s.c., minipump), and the other group (n=12) received CLB (in food pellets)+VPA (s.c., minipump). Video-EEG recordings were taken for another 14 days. EEG data and the videos were synchronized and written to a disk using custom software [11]. EEG recordings and video were stored in a custom file and MPEG4 formats. Seizures were defined as periods of high-frequency activity, 2x the baseline, lasting at least 10 seconds, followed by post-ictal depression. We calculated seizure frequency by dividing the number of seizures by the duration of recordings. Video-EEG reviewers were blinded to the treatment.

### Statistical analysis

All statistical analyses were conducted with GraphPad Prism 9.0. Data are presented as mean ± SD, and statistical significance was defined as a p-value<0.05. For the hyperthermia-induced test, a Log-rank (Mantel-Cox) test. A Wilcoxon test was used for paired non-parametric data, and a Mann-Whitney test was used for non-parametric data. A Fisher exact test was used to analyze contingency for categorical data.

## Results

### Subchronic treatment of STP with CLB and VPA was effective against hyperthermia-induced seizures in *Scn1a*^*A1783V/WT*^ mice

Before beginning efficacy studies for spontaneous seizures, it was necessary to determine a subchronic dosing strategy. PK data generated from acute injections of effective doses of STP add-on to CLB and VPA against a single hyperthermia-induced seizure test in *Scn1a*^*A1783V/WT*^ mice were used to calculate STP, CLB, and VPA sufficient to produce steady-state therapeutic concentrations when sub-chronically administered via drug-in-food pellets and osmotic minipump (VPA) [9]. In this study, we assessed whether a 14-day treatment of STP added to CLB and VPA would be effective against hyperthermia-induced seizures. STP (4000 mg/kg/day food pellet)+CLB (76 mg/kg/day food pellet)+VPA (1.5 g/mL at 0.5 μL/hr. flow rate) treatment resulted in a significant increase in threshold temperature to induce seizures on days 7 and 14 of treatment when compared to CLB (250 mg/kg food pellet)+VPA (1.5 g/mL at 0.5 μL/hr. flow rate) alone [median seizing temperatures: 41.1°C *p=0.0169 and 39.8°C **p=0.0014 respectively] or control pellet-treated *Scn1a*^*A1783V/WT*^ mice [median seizing temperature: 39.8°C *p=0.0232 respectively]. However, consistent with previous work from our lab, CLB+VPA treatment failed to increase the seizure threshold temperatures in *Scn1a*^*A1783V/WT*^ mice on Days 7 and 14 when compared to the control pellet treatment group [median seizing temperatures: 38.6°C p=0.1324 and 36.9°C; p=0.9781 respectively]. **(Figure 1)**. Furthermore, no signs of sedation or other adverse drug effects were observed in any animals in any group with this subchronic dosing strategy.

**Figure 1:**
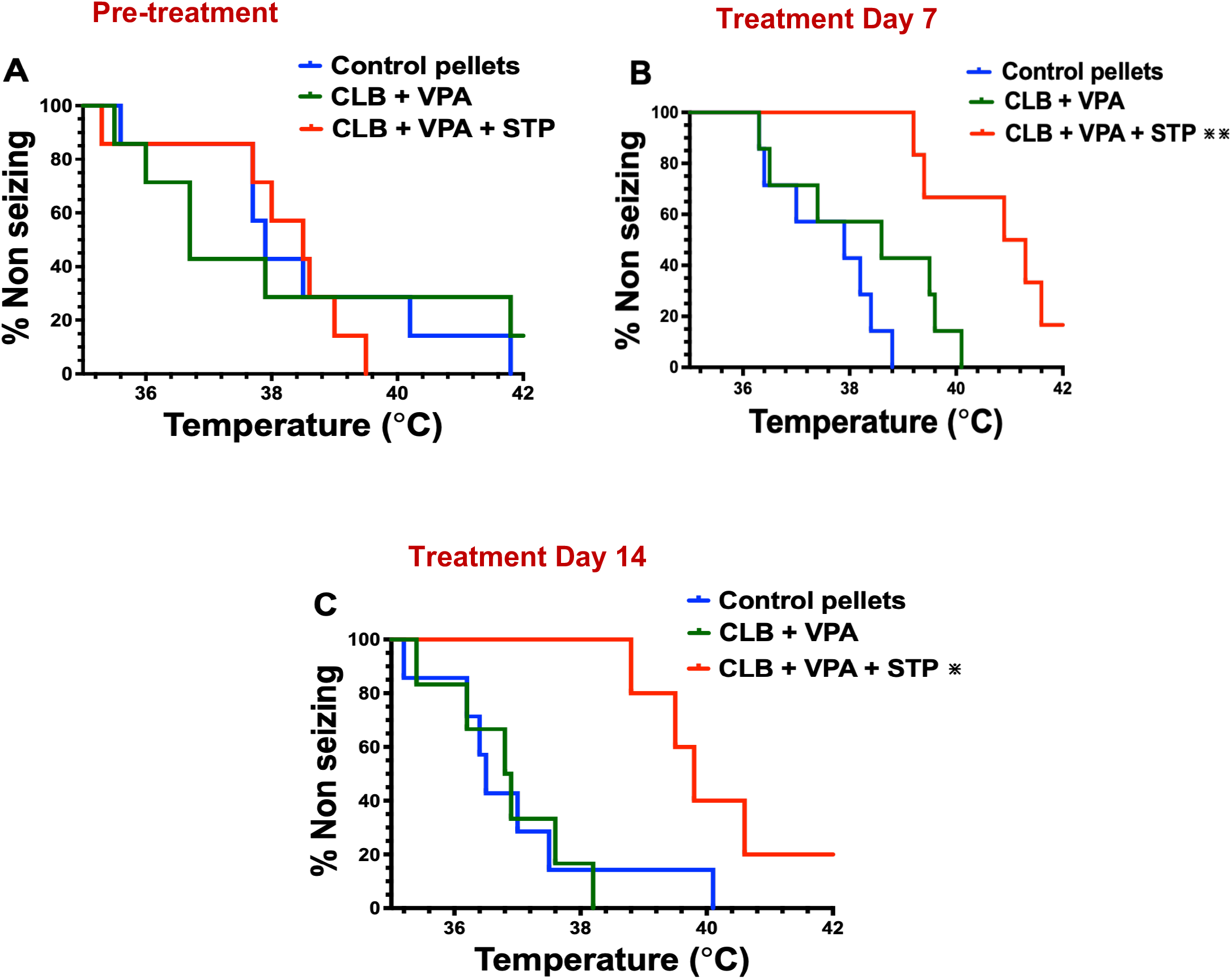
STP add-on to CLB and VPA significantly increased seizing temperature threshold after subchronic treatment in *Scn1a*^*A1783V/WT*^ mice. **(A)** At pretreatment (Day 0), mice in all 3 groups, control pellets (Bioserv Custom Diet + Saline) (n=7), CLB+ VPA (n=7), and CLB + VPA + STP (n=7) groups showed no significant differences in seizing temperature threshold. **(B)**. On treatment, Day 7, CLB + VPA + STP treatment significantly increased the temperature at which mice seized compared to the control pellet (n=7) andCLB + VPA (n=7) groups. **(C)**. On treatment, Day 14, CLB + VPA + STP continued to offer significant protection against hyperthermia-induced seizures compared to the control pellet group (n=7). At the same time, CLB + VPA (n=7) failed to increase the seizing temperature threshold in *Scn1a*^*A1783V/WT*^ mice. *p < .05, **p < 0.005; Log-rank (Mantel-Cox).

### Subchronic co-administration of CLB, VPA, and STP resulted in plasma concentrations within human therapeutic plasma concentration ranges

In the present study, we validate a preclinical animal model to help identify new ASMs that could be translated to the clinical setting in patients receiving polypharmacy. Thus, we assessed the predictive validity of our preclinical model and assay by determining if the pharmacokinetics of compounds in clinical use were like that seen in humans following subchronic treatment. We conducted bioanalytical studies to determine plasma and brain steady-state concentrations of STP, CLB, N-CLB, and VPA. Plasma and brain samples (n= 5-7) were taken from the same *Scn1a*^*A1783V/WT*^ mice cohort immediately after the hyperthermia-induced seizure efficacy test on Day 14 of treatment. All STP, CLB, and VPA doses resulted in plasma concentration within or slightly above the human plasma therapeutic range **(Table 1)**.

### STP add-on to CLB and VPA reduced spontaneous seizure burden in *Scn1a*^*A1783V/WT*^ mice

*Scn1a*^*A1783V/WT*^ mice have spontaneous seizures that can be measured with video-EEG **(Figure S1)**. We defined electrographic seizures as sharp-wave discharges with an increase in frequency and amplitudes at least twice the height of baseline, lasting longer than 10 seconds. Each electrographic ictal phase is synced with a video recording to match a behavioral seizure. Baseline video-EEG recordings were done for 2 weeks, followed by 2 weeks of drug treatment alongside video-EEG recording. We tested a combination of CLB and VPA against spontaneous seizures after subchronic administration, which did not offer significant protection against hyperthermia-induced seizures **(Figure 1)**. CLB (250 mg/kg/day food pellet)+VPA (1.5 g/mL at 0.5 μL/hr. flow rate) (n=12) did not significantly reduce the average number of SRS per mouse per day compared to baseline (p=0.0811**) (Figures 2A and 3A)**. However, STP (4000 mg/kg/day food pellet)+CLB (76 mg/kg/day food pellet)+VPA (1.5 g/mL at 0.5 μL/hr. flow rate) (n=13) significantly reduced the average number of SRS per mouse per day compared to baseline (*p=0.0327) **(Figures 2B and 3B)**. Moreover, STP add-on to CLB and VPA significantly reduced SRS frequency (***p=0.0003) compared to CLB+VPA (**Figure 2C)**. STP add-on to CLB and VPA also significantly increased seizure freedom during treatment in the *Scn1a*^*A1783V/WT*^ mice compared to CLB+VPA treatment, with 39% of the mice achieving seizure freedom (*p=0.0391), **(Figure 4)**.

**Figure 2:**
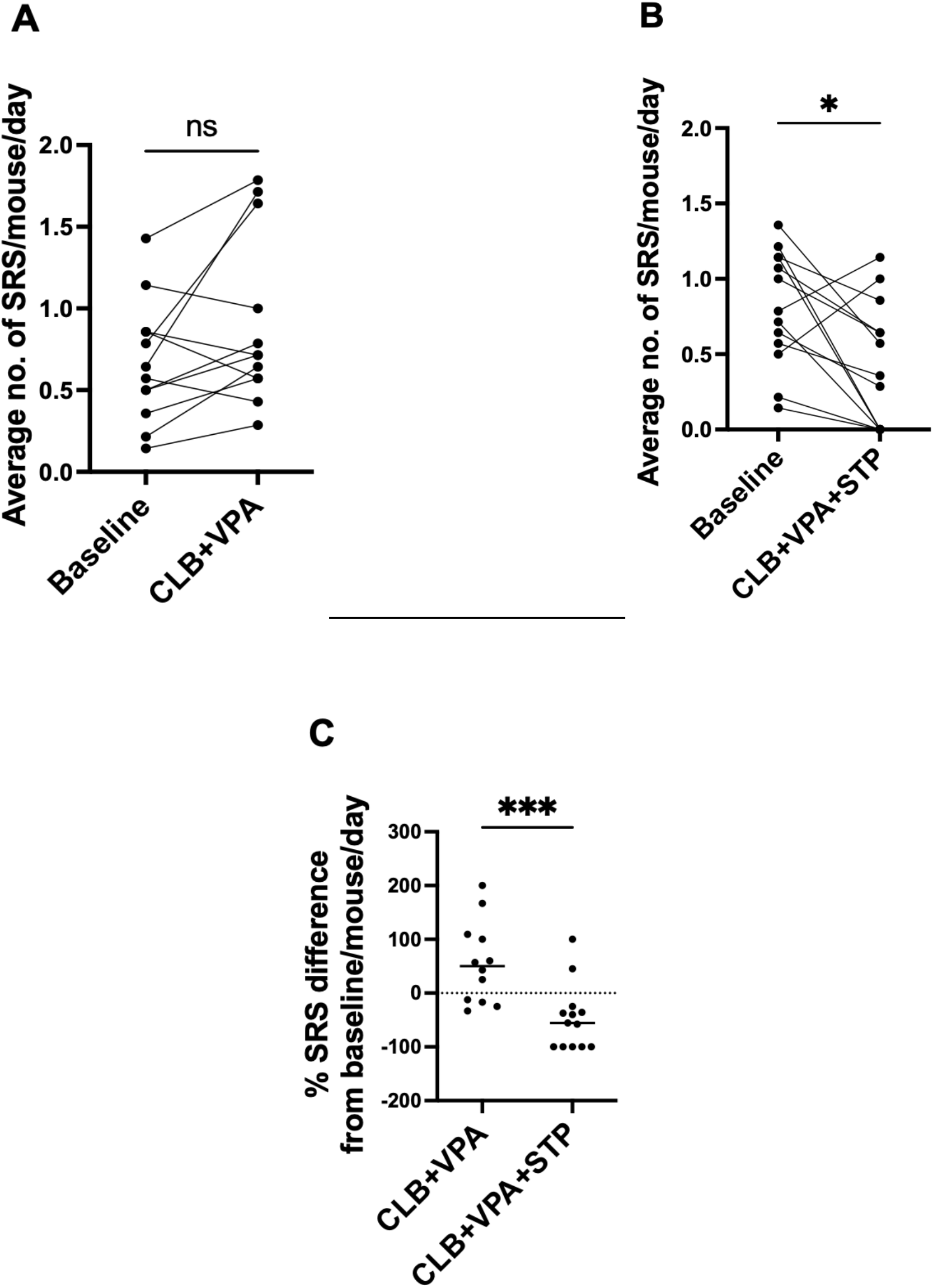
STP add-on to CLB and VPA significantly reduced the average number of seizures per day in *Scn1a*^*A1783V/WT*^ mice following subchronic treatment. **(A)** CLB + VPA drug treatment had no significant effect on seizure frequency when compared to baseline recordings over 14 days (n =12) p=0.0811, Wilcoxon Test. **(B)** CLB + VPA + STP drug treatment significantly reduced seizure frequency when compared to baseline recordings during a 14-day treatment period (n =13) *p=0.0327, Wilcoxon Test. **(C)** STP add-on to CLB + VPA (n=13) significantly reduced seizure frequency when compared to CLB + VPA (n=12) ***p=0.0003, Mann-Whitney Test.

**Figure 3:**
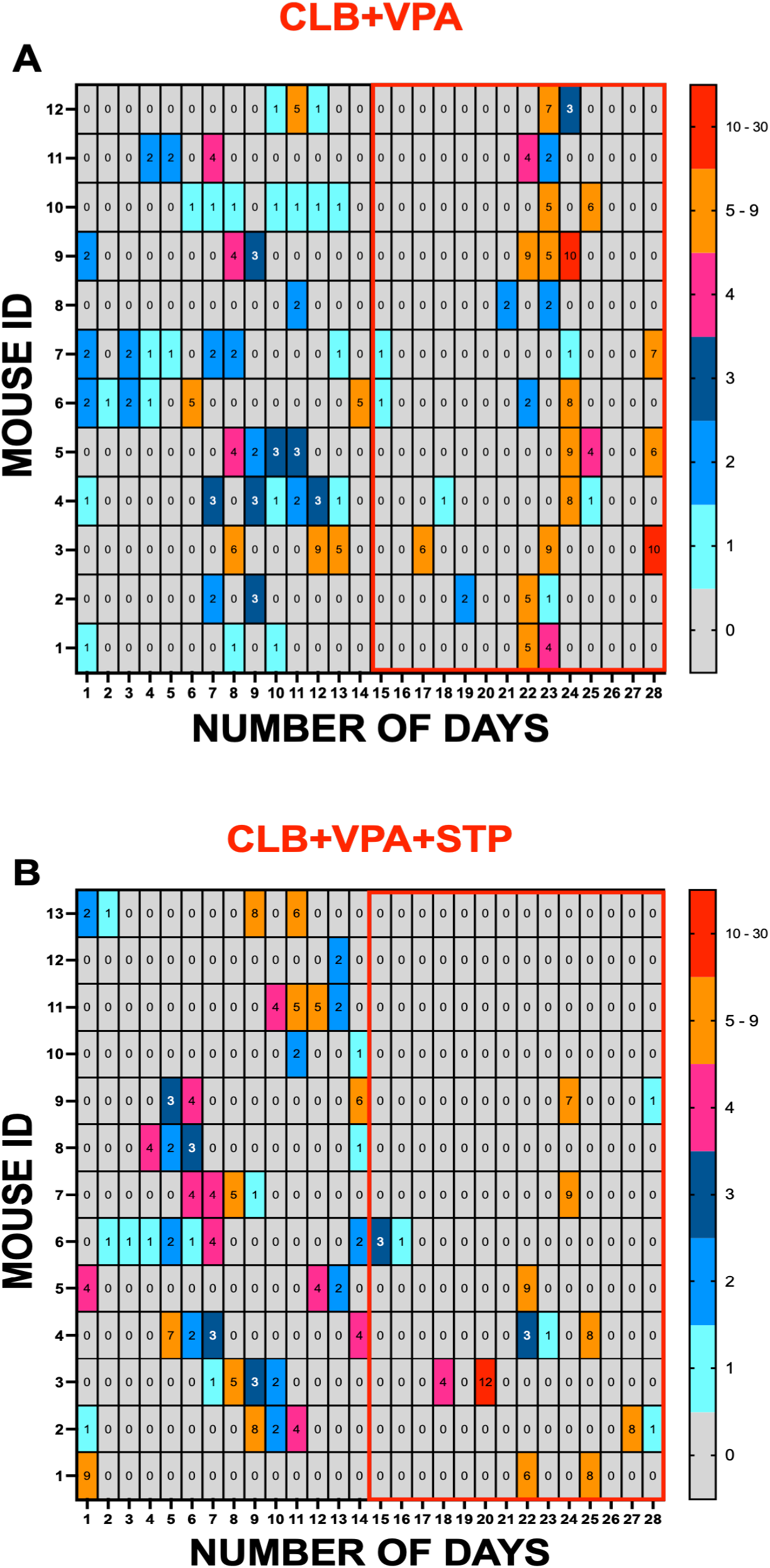
Heat Map of video-EEG analyses displaying SRS frequency pattern before and during treatment in *Scn1a*^*A1783V/WT*^ mice. **(A)** Heat map (gray to red) and number of seizures per day were recorded for each animal before (Day 1-14) and during CLB + VPA drug treatment (Day 15-28). **(B)** Heat map (gray to red) and number of seizures per day were recorded for each animal before (Day 1-14) and during CLB + VPA + STP drug treatment (Day 15-28). Treatment days are outlined in red.

**Figure 4:**
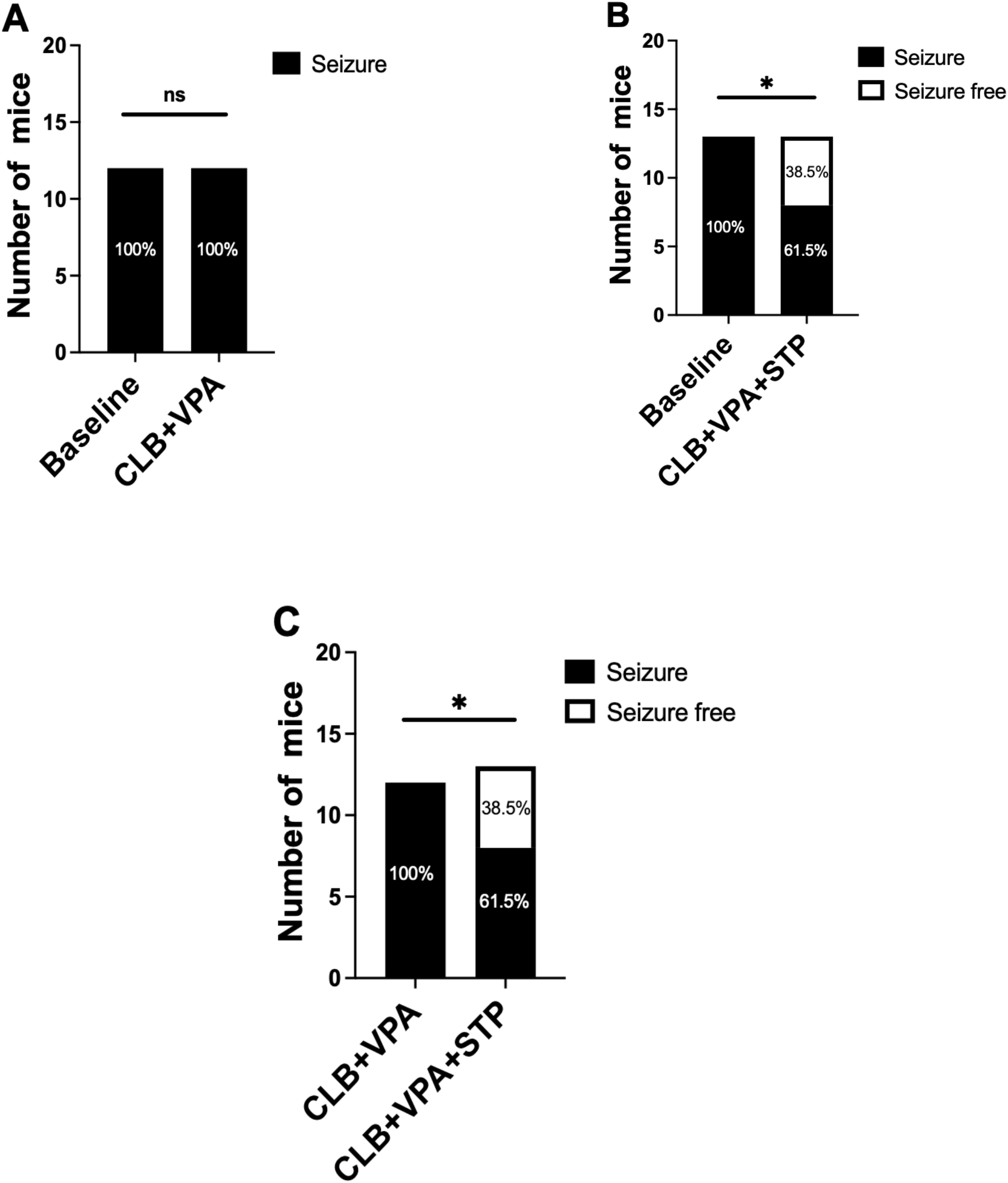
STP add-on to CLB and VPA significantly increased seizure freedom in *Scn1a*^*A1783V/WT*^ mice following subchronic treatment. **(A)** CLB + VPA drug treatment had no effect on seizure freedom when compared to baseline recordings over 14 days (n =12) p>0.999, Fisher exact Test. **(B)** CLB + VPA + STP drug treatment increased seizure freedom by 38.5% when compared to baseline recordings during a 14-day treatment period (n =13). **(C)** STP add-on to CLB + VPA (n=13) significantly reduced seizure frequency when compared to CLB + VPA (n=12); * p=0.0391, Fisher exact Test

## Discussion

Chronic monitoring of spontaneous seizures in rodent models of epilepsy is critical for preclinical epilepsy research [14]. Conventional preclinical development of efficacious ASMs has primarily relied on acute-induced seizure models [15, 16]. This shortfall in the preclinical approach may contribute to the significant number of pharmacoresistant patients with epilepsy [17-19]. Screening of subchronic drug dosing paradigm with continuous video-EEG monitoring in genetic models of epilepsy, such as the *Scn1a*^*A1783V/WT*^ DS mice, is technically challenging and resource-intensive [20]. However, this type of study may be necessary for successfully translating preclinical drug discovery into clinical use. Therefore, a drug screening approach that utilizes clinically relevant strategies would provide an efficient predictive measure for developing novel efficacious ASMs.

In the present study, we integrated PK evaluation of the ASMs to ensure that the dosage levels of the test drugs for the current study aligned with the human therapeutic plasma concentration range following a subchronic drug administration. The choice of drug administration strategy was guided by the PK data reported in a previous study from our lab [9]. We reported that CLB and STP have a half-life of ∼1.51 hr. when co-administered [9]; thus, in the current study, we administered both compounds via drug-in-food pellets using an automated feeder system that delivered six pellets at a 4-hour interval six times per day. Pellet consumption was tracked, and mice consumed all pellets delivered. Thus, this dosing approach proves advantageous by mitigating the stress experienced by the animal, which would occur with frequent drug administration via the i.p./s.c./oral gavage route over multiple days. However, due to the short half-life of VPA (∼0.6 hr.), we utilized the osmotic minipump, which delivers the drug subcutaneously at a constant rate of 0.5 μL/hr. The calculated doses with the administration routes for CLB, VPA, and STP used in this study achieved average steady-state concentrations within the human therapeutic plasma concentration range for either the parent compounds or their metabolites after a 14-day treatment in both the double (CLB + VPA) and triple-drug therapies (CLB + VPA +STP). A polytherapy approach that includes STP add-on to CLB and VPA has become the mainstay standard-of-care second-line treatment strategy for intractable seizures in people with DS [21, 22]. Similar to reports from several preclinical and clinical studies, we observed potential drug-drug interactions between CLB and its metabolite N-CLB and STP, resulting in elevated concentrations of CLB and N-CLB [9, 23-25]. Thus, a thorough understanding of the PK properties is crucial, especially in combination therapies, to facilitate the extrapolation of efficacy, safety, and tolerability in preclinical findings to predict clinical data [26, 27]. Inadequate considerations of preclinical PK assessment of ASMs during experimental design can result in misleading conclusions about a drug’s efficacy and safety, potentially leading to flawed extrapolations [28].

The present study also evaluated the efficacy of STP as an add-on therapy to CLB and VPA against hyperthermia-induced seizures using the subchronic dosing strategy informed by the PK studies. STP+CLB+VPA treatment significantly elevated the threshold temperature to induce seizures in *Scn1a*^*A1783V/WT*^ mice compared to CLB+VPA and vehicle groups on days 7 and 14 during a 14-day treatment regimen. Our data demonstrates that the efficacy of this triple-drug therapy was achieved at doses sufficient to provide steady-state therapeutic brain and plasma concentrations [29-31]. Similar to what was reported by Pernici et al. [8] with a single acute injection, coadministration of CLB with VPA was ineffective during a subchronic treatment regimen against hyperthermia-induced seizures in *Scn1a*^*A1783V/WT*^ mice. Similar to what is typically seen in clinical settings, not all patients are responsive to standard-of-care first-line therapies of CLB or VPA, either given alone or combined [32, 33]. This further suggests that hyperthermia-induced seizures in *Scn1a*^*A1783V/WT*^ mice are highly pharmacoresistant, and thus, this animal model could be very useful in identifying effective ASMs for drug-resistant seizures in persons with DS.

An often-cited limitation of preclinical drug screening using mouse models of epilepsy, particularly those involving genetic factors, is the use of only provoked seizure assays. In the present study, we further assessed the efficacy of a second-line treatment regimen, STP add-on to CLB and VPA, against SRS. STP+CLB+VPA treatment demonstrated seizure protective benefits against SRS in DS mice. This is consistent with reports from several randomized-control trials where greater than 53.3% responder rates were recorded when the efficacy of STP as an add-on to CLB and VPA was assessed [34, 35]. Complete seizure freedom is a desirable but challenging outcome to attain in highly drug-resistant epilepsies such as Dravet syndrome. Our data demonstrated that STP+CLB+VPA treatment increased seizure freedom compared to the CLB+VPA group. This is consistent with some clinical data reported from several clinical trials. Brigo et al. [36] reported that a significantly higher fraction of patients were seizure-free in the STP group compared with the placebo group. Moreover, Yildiz et al. [37] also noted that in a prospective, open-label, multicenter study conducted in Japan, ∼ 16% of DS patients achieved seizure freedom after a 56-week follow-up with STP treatment. STP add-on treatment achieved seizure freedom in 18% of patients with DS and non-DS refractory developmental and epileptic encephalopathies in a retrospective observational study [38]. We have demonstrated that the subchronic treatment approach of STP add-on to CLB and VPA therapy offers protection against febrile and spontaneous recurrent seizures in *Scn1a*^*A1783V/WT*^ mice. Thus, it would be essential for other FDA-approved and investigational add-on therapy approaches that are effective in increasing the temperature threshold to induce seizures to be further evaluated against spontaneous seizures when combined with CLB and VPA.

The current study has some limitations. Residual blood in the brain sample may impact measured brain concentrations since we did not perfuse brain tissue with saline before homogenization. The efficiency of the automated feeder system relied on the successful delivery of whole food pellets at the scheduled times and mice consuming the delivered pellets. Occasionally, we had feeders delivering crushed pellets, which could have influenced the total pellets consumed in a day and the drug dose administered.

Overall, our study demonstrated that subchronic treatment of STP combined with CLB and VPA offered protection against hyperthermia-induced and SRS frequency in *Scn1a*^*A1783V/WT*^ mice, similar to what is seen in clinical practice. While this was an expected outcome from these studies, it is essential to mention that the combined PK and efficacy approach described herein sets the stage for an effective screening platform to identify novel investigational ASMs with the potential to provide improved therapeutic outcomes for the person with drug-resistant epilepsy. Indeed, the present study establishes the framework for discovering novel and efficacious compounds. Future preclinical work may thus combine investigational compounds with CLB and VPA to determine if seizures can be adequately controlled. This screening approach is hypothesized to aid in identifying compounds that may prove efficacious in the person with DS.

## Acknowledgment

The authors thank the Epilepsy Therapy Screening Program at the National Institute of Neurological Disease and Stroke (NINDS) (ETSP) for their review and comments on this manuscript. The authors thank the staff at the Contract Site of the ETSP for their technical support.

## Supplementary Data

**Figure S1:**
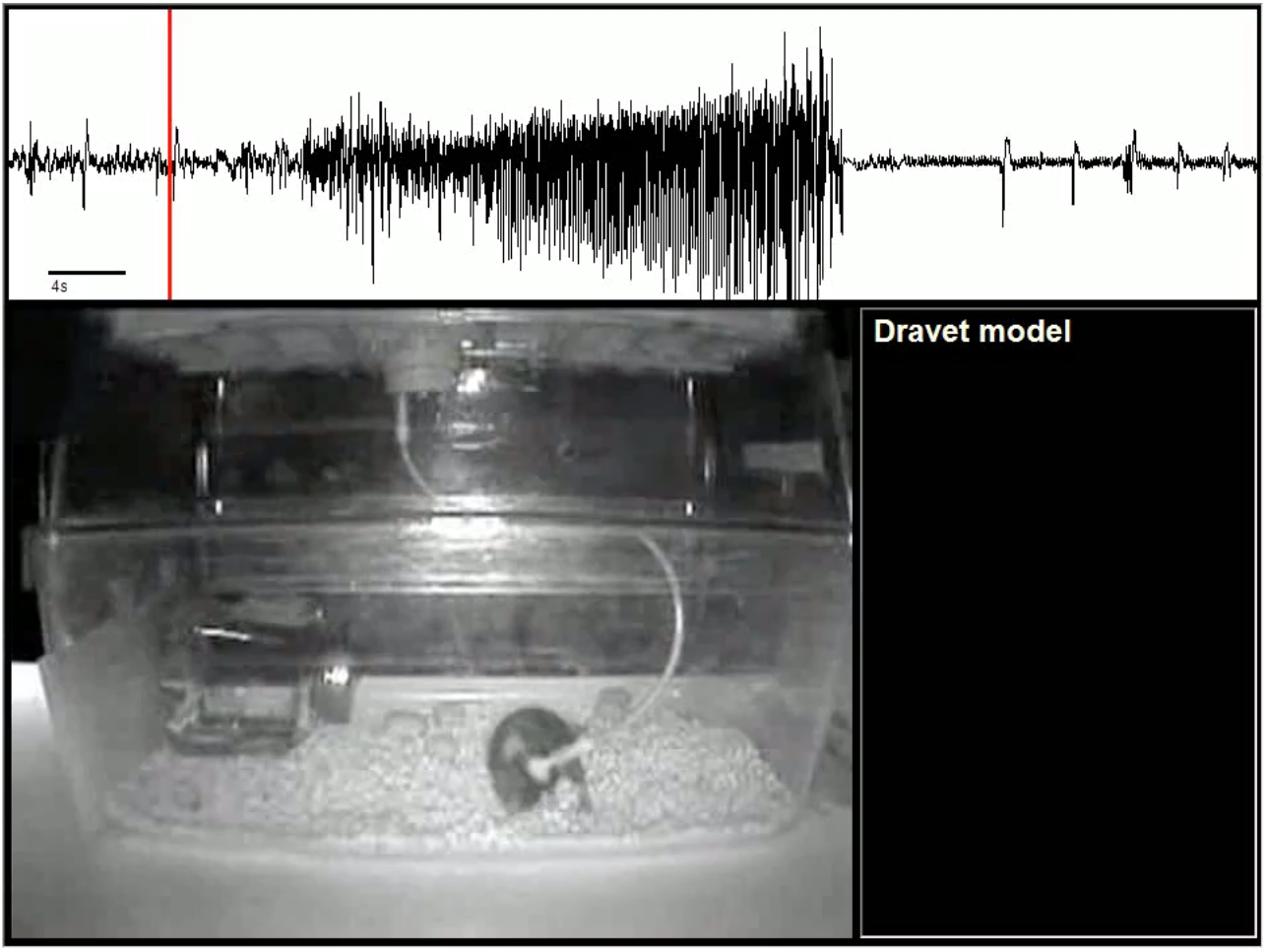
Video-EEG of a representative electrographic and behavioral spontaneous seizure.

## References

1. Wu, Y.W., Sullivan, J., McDaniel, S.S., Meisler, M.H., Walsh, E.M., Li, S.X., & Kuzniewicz, M.W. (2015). Incidence of Dravet Syndrome in a US Population. Pediatrics, 136(5), e1310–e1315. doi:10.1542/peds.2015-1807.

2. Brunklaus, A., Pérez-Palma, E., Ghanty, I., Xinge, J., Brilstra, E., Ceulemans, B., Chemaly, N., de Lange, I., Depienne, C., Guerrini, R., Mei, D., Møller, R., Nabbout, R., Regan, B., Schneider, A., Scheffer, I., Schoonjans, A., Symonds, J., Weckhuysen, S., Kattan, M.W., Zuberi, S.M. & Lal, D. (2022). Development and Validation of a Prediction Model for Early Diagnosis of SCN1A-Related Epilepsies. Neurology, 98(11), e1163–e1174. doi:10.1212/WNL.0000000000200028.

3. Chilcott, E., Díaz, J., Bertram, C., Berti, M., & Karda, R. (2022). Genetic therapeutic advancements for Dravet Syndrome. Epilepsy & Behavior, 132, 10874. doi: 10.1016/j.yebeh.2022.108741.

4. Kearney, J. (2013). Sudden unexpected death in Dravet Syndrome. Epilepsy Currents, 13(6), 264–265. doi:10.5698/1535-7597-13.6.264.

5. Wheless, J., Fulton, S., & Mudigoudar, B. (2020). Dravet syndrome: a review of current management. Pediatric Neurology, 107, 28–40. doi:10.1016/j.pediatrneurol.2020.01.005.

6. Griffin, A., Hamling, K., Hong, S., Anvar, M., Lee, L., & Baraban, S. (2018). Preclinical Animal Models for Dravet Syndrome: Seizure Phenotypes, Comorbidities and Drug Screening. Frontiers in Pharmacology, 9, 573. doi:10.3389/fphar.2018.00573.

7. Hawkins, N., Anderson, L., Gertler, T., Laux, L., George, A. J., & Kearney, J. (2017). Screening of conventional anticonvulsants in a genetic mouse model of epilepsy. Annals of Clinical and Translational Neurology, 4(5), 326–339. doi:10.1002/acn3.413.

8. Pernici, C., Mensah, J., Dahle, E., Johnson, K., Handy, L., Buxton, L., Smith, M., Peter, J., Metcalf, C., & Wilcox, K. (2021). Development of an Antiseizure Drug Screening Platform for Dravet Syndrome at the NINDS contract site for the Epilepsy Therapy Screening Program. Epilepsia, 62(7), 1665–1676. doi:10.1111/epi.16925.

9. Mensah, J., Johnson, K., Freeman, T., Reilly, C., Rower, J., Metcalf, C., & Wilcox, K. Utilizing an acute hyperthermia-induced seizure test and pharmacokinetic studies to establish optimal dosing regimens in a mouse model of Dravet syndrome. bioRxiv 2023.10.03.560653; doi:10.1101/2023.10.03.560653.

10. Wilcox, K., West, P., & Metcalf, C. (2020). The current approach of the Epilepsy Therapy Screening Program contract site for identifying improved therapies for the treatment of pharmacoresistant seizures in epilepsy. Neurpharmacolgy, 166, 107811. doi:10.1016/j.neuropharm.2019.107811.

11. Thomson, K., & White, H. (2014). A novel open-source drug-delivery system that allows for first-of-kind simulation of nonadherence to pharmacological interventions in animal disease models. Journal of Neuroscience Methods, 238, 105–111. doi:10.1016/j.jneumeth.2014.09.019.

12. Hawkins, N., Lewis, M., Hammond, R., Doherty, J., & Kearney, J. (2017). The synthetic neuroactive steroid SGE-516 reduces seizure burden and improves survival in a Dravet syndrome mouse model. Scientific Reports 7, 15327. 10.1038/s41598-017-15609-w.

13. Anderson, L., Hawkins, N., Thompson, C., Kearney, J., & George, A. J. (2017). Unexpected efficacy of a novel sodium channel modulator in Dravet syndrome. Scientific Reports, 10(7), 1682. doi:10.1038/s41598-017-01851-9.

14. Gu, B., & Dalton, K. (2017). Models and detection of spontaneous recurrent seizures in laboratory rodents. Zoological Research, 38(4), 171–179. doi:10.24272/j.issn.2095-8137.2017.042.

15. Löscher, W. (2011). Review: Critical review of current animal models of seizures and epilepsy used in the discovery and development of new antiepileptic drugs. Seizure, 20(5), 359–368. doi:10.1016/j.seizure.2011.01.003.

16. Simonato, M., Brooks-Kayal, A., Engel, J., Galanopoulou, A., Jensen, F., Moshé, S., O’Brien, T., Pitkānen, A., Wilcox, K., & French, J. (2014). The challenge and promise of anti-epileptic therapy development in animal models. Lancet Neurology, 13, 949–960. doi:10.1016/S1474-4422(14)70076-6.

17. Leclercq, K., Matagne, A., Provins, L., Klitgaard, H., & Kaminski, R. (2020). Pharmacological Profile of the Novel Antiepileptic Drug Candidate Padsevonil: Characterization in Rodent Seizure and Epilepsy Models. Journal of Pharmacology and Experimental Therapeutics, 372, 11–20. doi:10.1124/jpet.119.261222.

18. Kalilani, L., Sun, X., Pelgrims, B., Noack-Rink, M., & Villanueva, V. (2018). The epidemiology of drug-resistant epilepsy: a systematic review and meta-analysis. Epilepsia, 59, 2179–2193.

19. Kandratavicius, L., Balista, P.A., Lopes-Aguiar, C., Ruggiero, R.N., Umeoka, E.H., Garcia-Cairasco, N., Bueno-Junior, L.S., & Leite, J.P. (2014). Animal models of epilepsy: use and limitations. Neuropsychiatric Disease and Treatment, 10, 1693–1705. doi:10.2147/NDT.S50371.

20. Collard, R., Aziz, M., Rapp, K., Cutshall, C., Duyvesteyn, E., & Metcalf, C. (2022). Galanin analogs prevent mortality from seizure-induced respiratory arrest in mice. Frontiers in Neural Circuits, 6, 901334. doi:10.3389/fncir.2022.901334.

21. Cross, J., Caraballo, R., Nabbout, R., Vigevano, F., Guerrini, R., & Lagae, L. (2019). Dravet syndrome: Treatment options and management of prolonged seizures. Epilepsia, 60, S39–S48. doi:10.1111/epi.16334.

22. Wirrell, E., Laux, L., Donner, E., Jette, N., Knupp, K., Meskis, M., Miller, I., Sullivan, J., Welborn, M., & Berg, A. (2017). Optimizing the diagnosis and management of Dravet syndrome: Recommendations from a North American Consensus Panel. Pediatric Neurology, 68(18). doi:10.1016/j.pediatrneurol.2017.01.025.

23. Landmark, C., Johannessen, S., & Patsalos, P. (2020). Therapeutic drug monitoring of antiepileptic drugs: current status and future prospects,. Expert Opinion on Drug Metabolism & Toxicology, 16(3), 227–238. doi:10.1080/17425255.2020.1724956.

24. Giraud, C., Treluyer, J., Rey, E., Chiron, C., Vincent, J., Pons, G., & Tran, A. (2006). In vitro and in vivo inhibitory effect of stiripentol on clobazam metabolism. Drug Metabolism and Disposition, 34, 608–611. doi: 10.1124/dmd.105.007237.

25. Cardenal-Muñoz, E., Auvin, S., Villanueva, V., Cross, J., Zuberi, S., Lagae, L., & Aibar, J. (2022). Guidance on Dravet syndrome from infant to adult care: Road map for treatment planning in Europe. Epilepsia Open, 7, 11–26. doi: 10.1002/epi4.12569.

26. Brodie, M., & Sills, G. (2011). Combining antiepileptic drugs—rational polytherapy? Seizure, 20(5), 369–75. doi: 10.1016/j.seizure.2011.01.004.

27. Mahalmani, V., Sinha, S., Prakash, A., & Medhi, B. (2022). Translational research: Bridging the gap between preclinical and clinical research. Indian Journal of Pharmacology, 54(6), 393–396. doi:10.4103/ijp.ijp_860_22.

28. Kleiman, R., & Ehlers, M. (2016). Data gaps limit the translational potential of preclinical research. Science Translational Medicine, 8(320), 320ps321. doi:10.1126/scitranslmed.aac9888.

29. de Leon, J., Spina, E., & Diaz, F. (2013). Clobazam therapeutic drug monitoring: a comprehensive review of the literature with proposals to improve future studies. Therapeutic Drug Monitoring, 35(1), 30–47. doi:10.1097/FTD.0b013e31827ada88.

30. Methaneethorn, J. (2017). Population Pharmacokinetics of Valproic Acid in Patients with Mania: Implication for Individualized Dosing Regimens. Clinical Therapeutics, 39(6), 1171–1118. doi:10.1016/j.clinthera.2017.04.005.

31. Jacob, S., & Nair, A. (2016). An Updated Overview on Therapeutic Drug Monitoring of Recent Antiepileptic Drugs. Drugs in R D, 6(4). doi:10.1007/s40268-016-0148-6.

32. Lagae, L., Brambilla, I., Mingorance, A., Gibson, E., & Battersby, A. (2018). Quality of life and comorbidities associated with Dravet syndrome severity: a multinational cohort survey. Developmental Medicine and Child Neurology, 60(1), 63–72. doi:10.1111/dmcn.13591.

33. Wirrell, E., & Nabbout, R. (2019). Recent advances in the drug treatment of Dravet syndrome. CNS Drugs, 33(9), 867–881. doi:10.1007/s40263-019-00666-8.

34. Brigo, F., Igwe, S., & Bragazzi, N. (2020). Stiripentol add-on therapy for drug-resistant focal epilepsy. Cochrane Database Syst Rev, 5(5), CD009887. doi:10.1002/14651858.CD009887.pub5.

35. Janković, S., Janković, S.V., Vojinović, R., & Lukić, S. (2023). Investigational new drugs for the treatment of Dravet syndrome: an update. Expert Opinion on Investigational Drugs, 32(4), 325–331. doi:10.1080/13543784.2023.2193680.

36. Brigo, F., Igwe, S., & Bragazzi, N. (2017). Antiepileptic drugs for the treatment of infants with severe myoclonic epilepsy. Cochrane Database. Syst Rev., 5(:Cd010483.).

37. Yildiz, E., Ozkan, M., Uzunhan, T., Bektaş, G., Tatlı, B., Aydinli, N., Çalışkan, M., & Özmen, M. (2019). Efficacy of Stiripentol and the Clinical Outcome in Dravet Syndrome. Journal of Child Neurology, 34(1), 33–37. doi:10.1177/0883073818811538.

38. Gil-Nagel, A., Aledo-Serrano, A., Beltrán-Corbellini, Á., Martínez-Vicente, L., Jimenez-Huete, A., Toledano-Delgado, R., Gacía-Morales, I., & Valls-Carbó, A. (2024). Efficacy and tolerability of add-on stiripentol in real-world clinical practice: An observational study in Dravet syndrome and non-Dravet developmental and epileptic encephalopathies. Epilepsia Open, 9(1), 164–175. doi:10.1002/epi4.12847.

